# PLAMseq enables the proteo-genomic characterization of chromatin-associated proteins and protein interactions in a single experimental workflow

**DOI:** 10.1101/2025.04.27.650851

**Authors:** Lourdes González-Vinceiro, Carmen Espejo-Serrano, María Eugenia Soler-Oliva, María Luisa Mateos-Martín, Daniel Rico, Cristina González-Aguilera, Román González-Prieto

**Affiliations:** Andalusian Centre for Regenerative Medicine and Molecular Biology (CABIMER), Universidad de Sevilla-CSIC-Universidad Pablo de Olavide-Junta de Andalucía, Sevilla, Spain; Departamento de Biología Celular, Facultad de Biología, Universidad de Sevilla, Sevilla, Spain; Institute of Biomedicine of Seville, IBiS/Hospital Universitario Virgen del Rocío/CSIC/Universidad de Sevilla, Proteomics Facility, Sevilla, Spain

## Abstract

Chromatin Immunoprecipitation (ChIP) and Co-Immunoprecipitation (CoIP) assays are common approaches to characterize the genomic localization and protein interactors, respectively, for a protein of interest. However, these approaches require the use of specific antibodies, which often face sensitivity and specificity issues. Based on TurboID, we developed PLAMseq (Proximity Labelled Affinity-purified Mass spectrometry plus sequencing), which enables, in the same workflow, to identify the genomic loci and the interacting proteome of a protein of interest. Moreover, PLAMseq can also be applied to specifically map protein interactions and ubiquitin(-like) modified proteins.

We validated PLAMseq with two well characterized proteins, RNA polymerase II and CTCF, with excellent robustness and reproducibility. Next, we applied PLAMseq to characterize Histone H1 SUMOylation, which study has remained elusive due to the lack of specific reagents, and found that SETDB1 binds to SUMOylated histone H1.2 and H1.4 which also colocalize with H3K9me3 at repetitive regions of the genome.

## INTRODUCTION

Our genomes are scaffolded onto a protein structure conforming the chromatin, which main elemental unit is the nucleosome, consisting of one pair of each of the core histones, H2A, H2B, H3 and H4 and the linker histone H1. Histone Post Translational Modifications (PTMs) are the main source of epigenetic information. The functional study of chromatin factors and modifications requires the determination of their distribution across the genome and the identification of partners and components of functional complexes.

To study the distribution of proteins across the genome, several approaches have been developed, including Chromatin ImmunoPrecipitation and sequencing (ChIPseq) ^2^, Cut and Run ^3^ and Cut and Tag ^4^, all of them requiring specific high sensitivity antibodies (i.e. ChIP grade) for their application. Alternatively, DamID is an antibody-free approach ^5^ which tags a protein of interest with the dam methylase making the interacting DNA sensitive to dam-specific restriction enzymes. DamID has a resolution limitation compared to the other approaches ^6,7^.

For the identification of protein interactors, mass spectrometry-based proteomics approaches coupled to different enrichment strategies are available. Co-immunoprecipitation (CoIP) experiments have been widely applied. However, detecting transient interactions by CoIP is very challenging. More recently, this challenge has been addressed by Proximity-labelling strategies based on the biotinylation of the proximal proteome. First approach was BioID, consisting of tagging the protein of interest with a promiscuous mutant of *E*.*coli* biotin-ligase BirA ^8^. BioID had a few drawbacks, first BirA had affinity for the DNA, and it needed hours to label its proximity proteome. The affinity for DNA problem was solved with BioID2 using a mutant biotin-ligase from *Aquifex aeolicus* ^9^. To solve the need of long labelling times, a much faster approach for biotin labelling is based in ascorbate peroxidase and using biotin-phenol as substrate upon the addition of H_2_O_2_. This approach is termed APEX, which was later improved as APEX2 ^10,11^. APEX can label proximal proteins within minutes. The main problem of APEX(2) is that it requires treating the cells with H_2_O_2,_ which can be very toxic.

More recently, an improved version of BioID, TurboID, achieved a rapid labelling without toxicity ^12^. Several adaptations of TurboID have been described to study chromatin related processes such as Protein A-TurboID ^13^. Protein A-TurboID enables the identification of the proximal proteome of a specific protein or PTM-modified protein without the need of tagging the protein of interest. It uses a specific antibody that is then recognized by a Protein A-TurboID fusion to catalyze the biotinylation reaction. Another adaptation is TurboCas ^14^, which fuses non-catalytic Cas9 to TurboID. Turbo-Cas can be directed to specific genomic loci using a gRNA and then biotinylate the proteome associated to such locus.

Here we report PLAMseq (<u>P</u>roximity <u>L</u>abelled <u>A</u>ffinity-purified <u>M</u>ass spectrometry and <u>seq</u>uencing) a new application of TurboID to perform proteomics and genomics simultaneously, in the same workflow. Given that the biotin-streptavidin is the most stable interaction described in nature, we hypothesized that biotinylated proteins from TurboID would resist the crosslinking-decrosslinking processes performed in ChIP-seq workflows and remain attached to the streptavidin. Indeed, we show that it is possible to induce the activity of TurboID by the addition of biotin, shortly after induce DNA-protein crosslinking and then co-purify the proteins with its associated DNA using streptavidin beads. Next, Protein-DNA crosslinks are reverted and the DNA sequenced, subsequently the proteins are trypsin-digested and identified by mass spectrometry-based proteomics (Figure 1). Moreover, TurboID allows Bimolecular Complementation (BiC), as split-TurboID ^12^ making this approach also applicable to study protein complexes and interactions in their genomic context.

**Figure 1.**
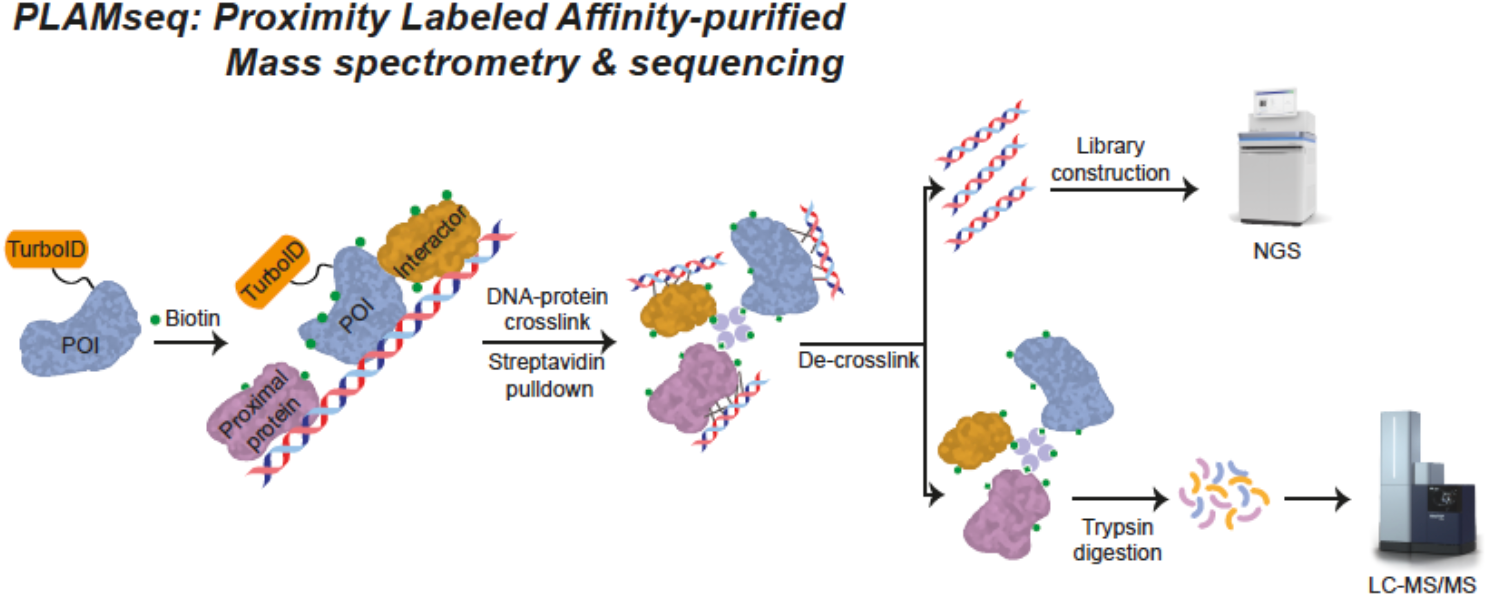
Cartoon depicting the PLAMseq approach. After as short biotin pulse, proteins proximal to the TurboID-tagged protein of interest become biotinylated and crosslinked to DNA with formaldehyde. Proteins and DNA are co-purified using Streptavidin beads. Next DNA is de-crosslinked, purified and sequenced. Subsequently, proteins are trypsin-digested and analyzed by mass spectrometry-based proteomics.

## RESULTS

### PLAMseq of CTCF and RNApol II

In order to test the validity and performance of our PLAMseq approach, we applied it with two proteins which localization in the genome is very well defined. Namely, CTCF, a protein involved in genomic loop extrusion ^15-17^, and which genomic loci have been previously defined, and RNA polymerase II.

First, we constructed a lentiviral plasmid for the systematic stable-inducible expression of TurboID-tagged proteins. This plasmid contained a 10xHIS + FLAG tag followed by Gateway® cloning cassette, a GGS linker and TurboID protein which enabled the straightforward shuttling of any gene of interest from a Gateway® donor or entry plasmid.

Then, we cloned CTCF in our doxycycline-inducible TurboID plasmid. Next, we introduced the inducible CTCF-TurboID construct in HeLa cells by lentiviral transduction and tittered the expression level to express it at near-to-endogenous levels with different doxycycline concentrations (Supplementary Figure 1A). We observed a reduction of the endogenous CTCF levels proportional to the expression of the CTCF-TurboID construct. Thus, the total CTCF levels after induction were virtually endogenous.

We induced the expression of CTCF-TurboID at 1 µg/ml and performed PLAMseq in three independent biological replicates. After applying a 10-minute biotin-pulse, we fixed the cells by inducing DNA-protein crosslinks with formaldehyde. Next, cells were lysed, DNA fragmented by sonication and TurboID-labelled proteins and their associated DNA molecules purified with Streptavidin beads. Subsequently, DNA-Protein crosslinks were reverted releasing the DNA, which was purified and processed for Illumina sequencing, while the proteins kept retained on the Streptavidin beads. Finally, proteins were trypsin-digested and analyzed by mass spectrometry-based proteomics. Cells not expressing the TurboID constructs were used as negative control.

Analysis of the CTCF proximal proteome revealed, as expected, CTCF as the top hit (Figure 2A, Supplementary Figure 1B, Supplementary Dataset 1). Moreover, other proteins previously described to interact with CTCF were also identified, such as SMARCA5 ^18^, ZNF280C ^19^, or the Cohesin complex ^20,21^. Furthermore, genomics analysis (Figure 2B-C) showed that 24855 significant CTCF PLAMseq peaks were identified above the Control sample and 85% of them localized at previously annotated CTCF sites in the HeLa S3 genome obtained by ChIP-seq (Encode ChIP-seq). These results validate the applicability of PLAMseq approach for proteo-genomic characterization of chromatin-associated proteins. Importantly, heatmaps of CTCF signal around the encode CTCF sites revealed that PLAMseq identified narrower peaks compared to ChIP-seq indicating that a more precise identification of CTCF binding in the genome can be obtained in PLAMseq (Figure 2D).

**Figure 2.**
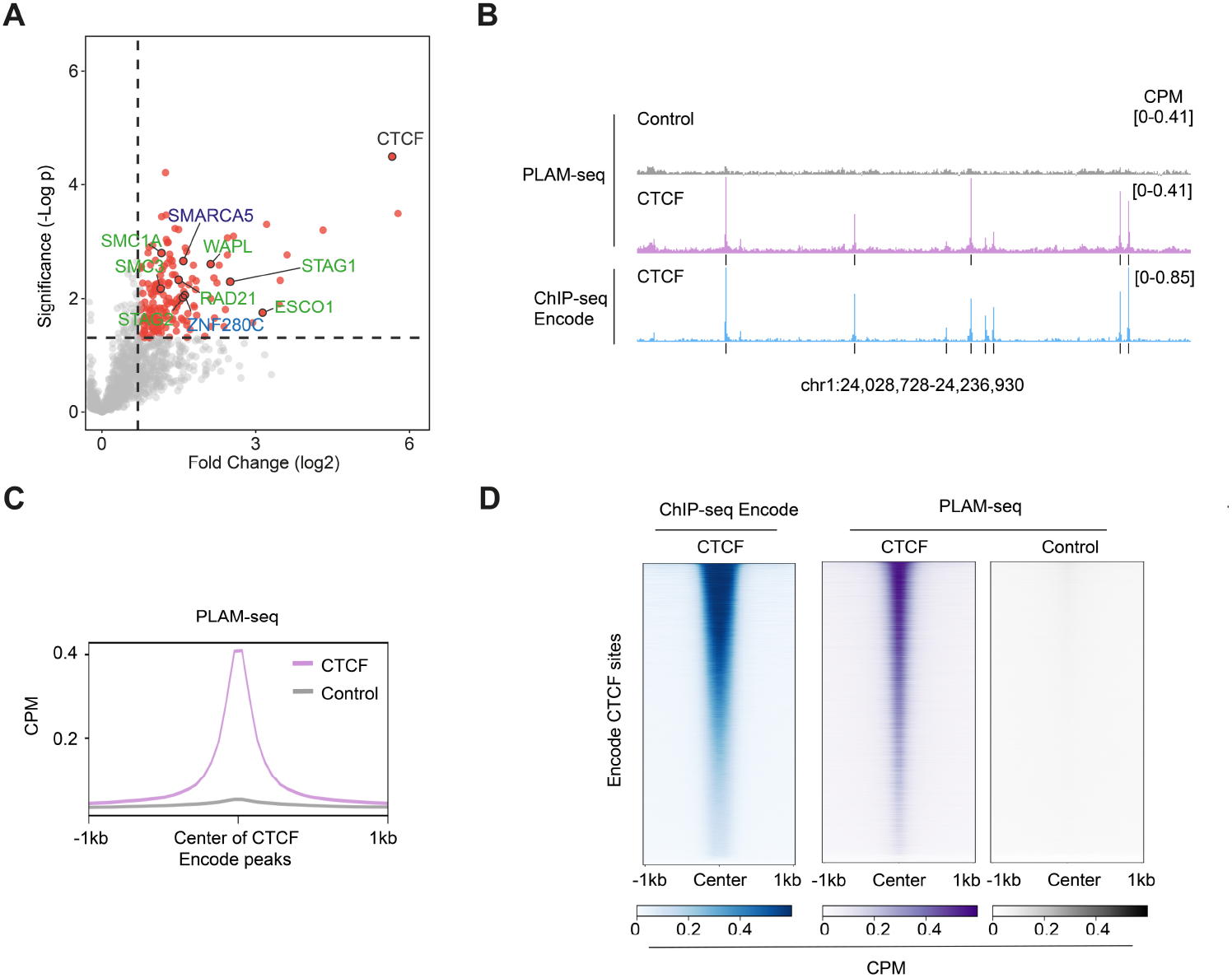
**(A)** Volcano Plot depicting statistical differences in proteomics analysis of PLAMseq samples comparing CTCF to POLR2I. Only the CTCF side of the Volcano is shown. Each dot represents a protein and selected proteins are labelled. Proteins related to the same complex are color matched. Dotted lines mark significance threshold for p>0.05 from unpaired two-tailed t-tests and fold change higher than 0.7 (log2). **(B)** Maps of CTCF and control samples identified by PLAMseq or ChIPseq. Black boxes bellow CTFC signal represent significant peaks where CTCF binds. **(C)**. Average profiles of PLAMseq CTCF and Control signal plotted around 2kb of the center of CTCF binding sites identified in ChIPseq samples from ENCODE. **(D)**. Heatmaps showing the ChIPseq signal or the PLAMseq signal from CTCF or Control samples around CTCF binding sites identified in ChIPseq samples from ENCODE. In all samples, CTCF sites were sorted from maximum to minimal intensity found in ChIPseq from ENCODE.

Nevertheless, CTCF has a very discrete distribution across the genome at very well-defined sites. Thus, we decided to investigate the validity and performance of the PLAMseq approach for proteins with a wider genomic distribution. Therefore, we applied PLAMseq, to investigate the genomic distribution and proximity proteome of RNA polymerase II. RNA polymerase II is a complex made of several proteins, including the POLR2I subunit. We made a POLR2I-TurboID construct (Figure 3, Supplementary Figure 1C). Mass spectrometry-based proteomics analysis of the POLR2I proximity proteome (Figure 3A, Supplementary Figure 1D, Supplementary Dataset 1) reveled many subunits of RNA Polymerase II, indicating that the POLR2I-TurboID construct is incorporated in the RNA Polymerase II complex. Additionally, we identified other RNA Polymerase II-associated proteins such as PAF1 and LEO1^22^, the integrator complex INTS1 ^23^, NELF complex ^24^ or PHF3 ^25^. Genomics analysis shows that POLR2I PLAMseq samples have a better signal-to-noise ratio than the published RNA Polymerase II ChIP-seq (ENCODE) (Figure 3B-C) and that, similar to published data, signal mainly accumulated at the promoter regions of transcribed genes, with a decreasing trend along the gene body (Figure 3B-C). Consistently, 84.3% of RNAPII peaks localize within the 500bp upstream of annotated Transcription Start Sites (TSS). We also demonstrated that the PLAMseq binding is specific for actively transcribed genes since signal in silent genes is dramatically reduced (Figure 3D). Altogether these data demonstrate that PLAMseq can efficiently identify protein interactors and DNA binding sites of a protein of interest in a unified protocol.

**Figure 3.**
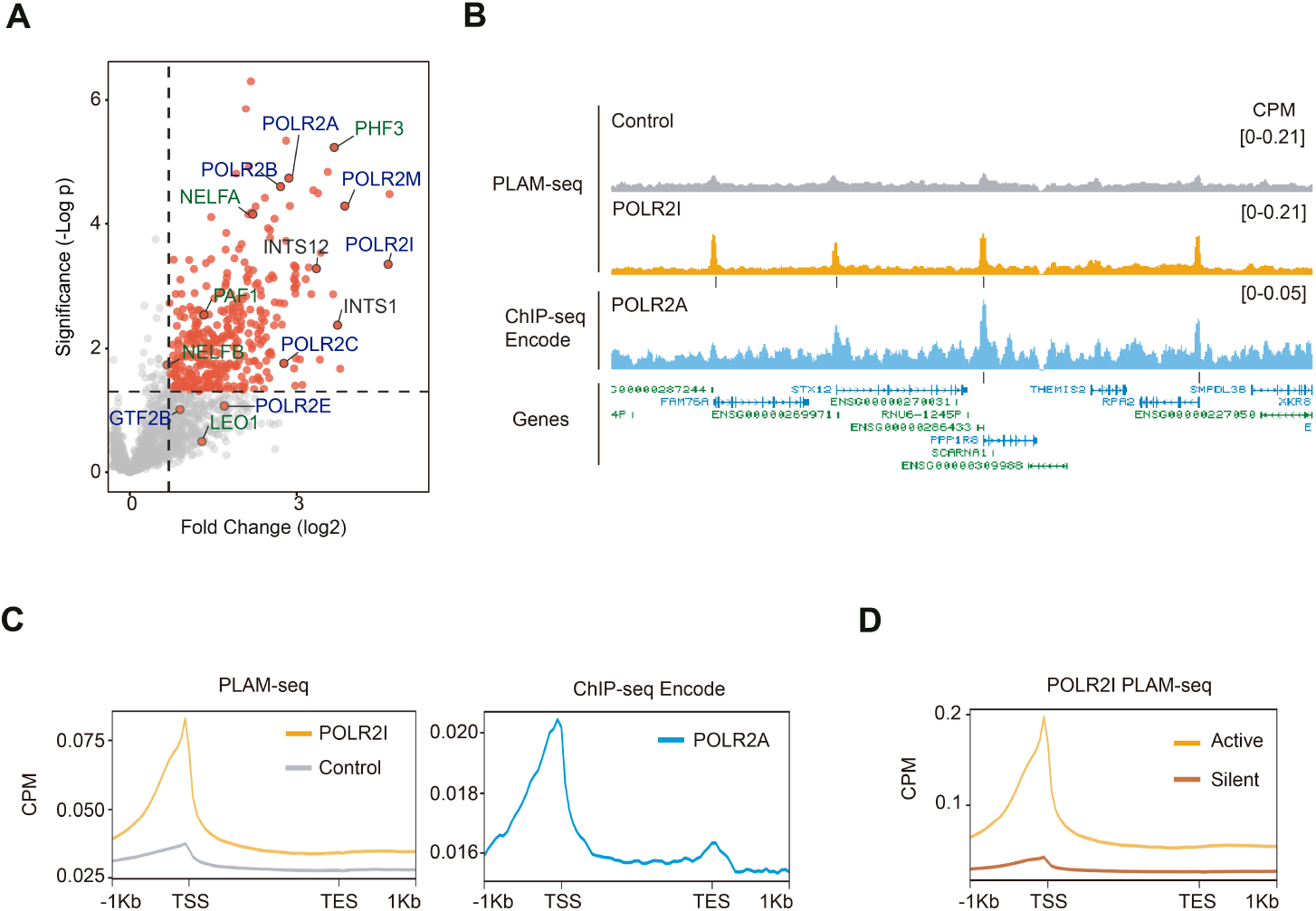
**(A)** Volcano Plot depicting statistical differences in proteomics analysis of PLAMseq samples comparing POLR2I to CTCF. Only the POLR2I side of the Volcano is shown. Each dot represents a protein and selected proteins are labelled. Proteins related to the same complex are colour-matched. Dotted lines mark significance threshold for p>0.05 from unpaired two-tailed t-tests and fold change higher than 0.7 (log2). **(B)** Maps of POLR2I, control and POLR2A samples identified by PLAMseq or ChIPseq, respectively. Black boxes bellow RNAPII signal represent significant peaks where these RNAPII subunits bind. **(C)** Average profiles of PLAMseq (left) and ChIPseq (right) signals from samples described above and plotted around all genes. Genes were scaled to 2 kilobases (Kb) to correct for gene length differences. **(D)** Average profiles of PLAMseq of POLR2I plotted around transcriptionally active and silent genes. Genes were scaled as in C.

Reproducibility of sequencing results for both CTCF and POLR2I was high as shown in the high Pearson correlation between replicates in sequencing results (Supplemental Figure 1E).

### Using PLAMseq to study Histone SUMOylation

One of the advantages of TurboID is that it allows Bimolecular Complementation (BiC), also known as split-TurboID ^12^. In the split-TurboID version, two proteins are tagged each with one complementary fragment of the TurboID enzyme. When these two proteins interact or are in close proximity, the TurboID enzymatic activity is reconstituted and proteins proximal to this interaction become biotin-labelled.

Using split-TurboID, the Rosa Barrio’s laboratory recently published SUMO-ID, a method to identify SUMO-dependent interactors of specific proteins ^1^. Among the chromatin-related proteins, histones conform the elemental chromatin unit, the nucleosome. All histone types have been described to be SUMOylated in several high-throughput mass spectrometry-based proteomics screens ^26-29^.

However, our understanding of the functional relevance of histone SUMOylation is very limited. Studies in yeast indicated that SUMO conjugation on histone H4 lysine 12 (H4K12Su) is associated with chromatin decompaction and repression of transcription mediated by the p300 histone acetylating complex ^30-32^. In contrast, other studies in mouse embryonic stem cells showed that histone H1 SUMOylation promotes chromatin condensation and restricts embryonic cell fate identity ^33^. In both cases no mechanistical model has been proposed about the function of histone H1 SUMOylation.

Therefore, we decided to adapt SUMO-ID to our PLAMseq approach (Figure 4A) to characterize histone H1 SUMOylation in a proteo-genomic manner. As a model, we used HeLa cells, as Histone H1 has been previously described to be SUMOylated in this cell line ^28^. As we were uncertain about how stable histone H1 SUMOylation is, we made some modifications on the original myc-CTurboID-SUMO constructs for SUMO1 and SUMO2. First, we reverted the SUMO1^Q95P^ and SUMO2^Q90P^ mutations to make the constructs cleavable again by SUMO proteases. Additionally, we introduced the SUMO1^Q92R^ and SUMO2^Q88R^ mutations, so the SUMO tryptic remnants enable the identification of SUMOylation sites by mass spectrometry-based proteomics ^34,35^. Finally, we introduced them in stable expression lentiviral backbones and transduced each of them separately into HeLa cells.

**Figure 4.**
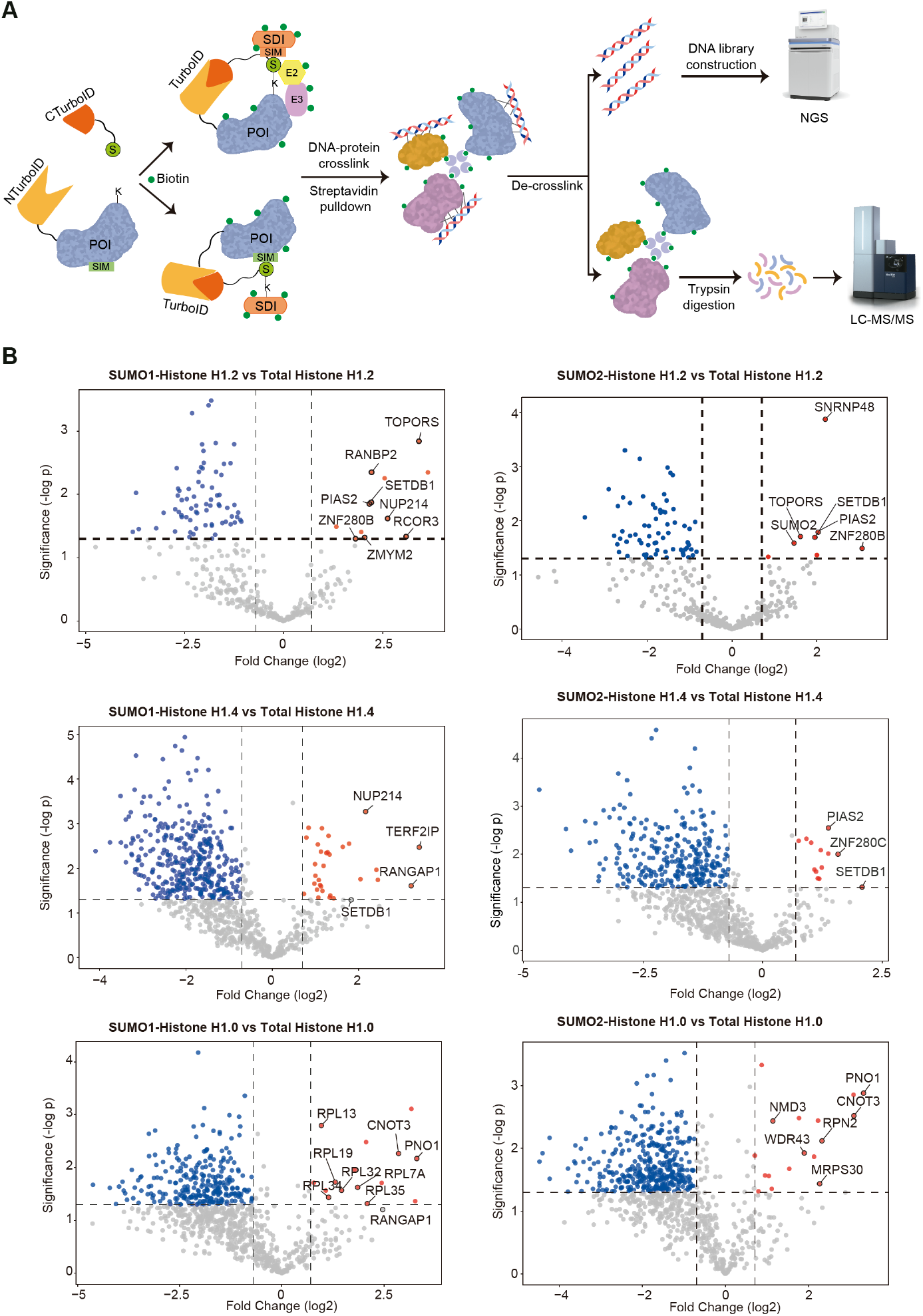
**(A)** Cartoon depicting the split-PLAMseq approach with SUMO-ID. SUMO is tagged with the C-terminal half of TurboID and the Protein of Interest is tagged with the N-terminal half of Turbo-ID. After as short biotin pulse, proteins proximal to the SUMOylated protein of interest become biotinylated and crosslinked to DNA with formaldehyde. Proteins and DNA are co-purified using Streptavidin beads. Next DNA is de-crosslinked, purified and sequenced. Subsequently, proteins are trypsin-digested and analyzed by mass spectrometry-based proteomics. The SUMO-ID carton is adapted with modifications from ^1^. **(B)** Volcano plots depicting statistical differences from PLAMseq proteomics samples between SUMOylated histone H1 types with either SUMO1 or SUMO2 with their respective total histone H1 counterparts. Each dot represents a protein and selected proteins are labelled. Dotted lines mark significance threshold for p>0.05 from unpaired two-tailed t-tests and fold change higher than 0.7 (log2).

The new myc-CTurboID-SUMO constructs were expressed in a stable manner and were able to form conjugates with acceptor proteins while total SUMO levels remained close to endogenous (Supplementary Figure 2A-B). Next, we introduced different NTurboID-Histone H1 types by lentiviral transduction. To be able to compare SUMOylated Histone H1 with total histone H1, we introduced the same histone H1 types tagged with full length TurboID.

Among the different histone H1 types, H1.2 is the most highly SUMOylated type in HeLa cells, followed by H1.4 ^28^. Therefore, we decided to study these two types of histone H1. Additionally, we included in our study the non-replicative histone H1.0, which sumoylation has been observed after proteasome inhibition in HeLa cells ^28^, and more recently in normal conditions in mouse cells ^29^. In summary, we performed comparative analysis of PLAMseq and SUMO-ID-PLAMseq samples of histones H1.2, H1.4, and H1.0.

Proteomics analysis (Figure 4B, Supplementary Datasets 2-4) showed that SUMOylation-related proteins preferentially interacted with SUMOylated histone H1.2 (compared with total histone H1.2). These include the SUMO E3 enzyme PIAS2, which can act both with SUMO1 and SUMO2 ^36^, the SUMO1-Targeted Ubiquitin Ligase (STUbL) TOPORS ^37^ and the SUMO1-specific E3 RANBP2 for SUMO1-H1.2.

Interestingly, SETDB1 was identified as a SUMOylated histone H1.2 preferential interactor, both for SUMO1 and SUMO2. SETDB1 is the most important methyltransferase for H3K9me3 ^38^ SETDB1 was also a preferential interactor for SUMOylated histone H1.4. In contrast, in the case of SUMOylated histone H1.0, the most enriched interactors were mainly components of ribosomal particles.

In the PLAMseq genomics analysis, we observed that the genome-wide profiles of the different SUMOylated histone H1 types show a similar genomic distribution than their total histone H1 counterpart (Supplementary Figure 3). Nevertheless, proteomic analysis had identified for H1.2 and H1.4 a SUMO-preferential interaction for SETDB1. SETDB1 is the main writer of the H3K9me3 histone mark, which defines constitutive heterochromatin at repetitive elements. However, these elements are not well represented in the GRCh38/hg38 reference genome we had used to align our histone H1 and SUMO-histone H1 PLAMseq data. Therefore, we decided to re-align our PLAMseq data to the Telomere-to-Telomere T2T-CHM13 reference genome based on long reads ^39^.

To functionally annotate the T2T-CHM13-aligned HeLa epigenome, we also re-aligned publically available histone PTM ChIP-seq datasets and used them to define 11 different chromatin states. The combinatorial presence of histone PTMs in these states are characteristic of different functional genomic features ^40^. Namely, H3K27me3 and H3K9me3 mark facultative and constitutive heterochromatin, respectively, while H3K36me3, H3K4me1, H3K27ac or H3K4me3 highlight different euchromatin features (Figure 5A) ^38^. Next, we investigated how the relative enrichment of SUMOylated Histone H1 types compared to their total H1 counterparts localized in these chromatin states (Figure 5B). Additionally, the CTCF and POLR2I PLAMseq data was included in the analysis together with other genomic features.

**Figure 5.**
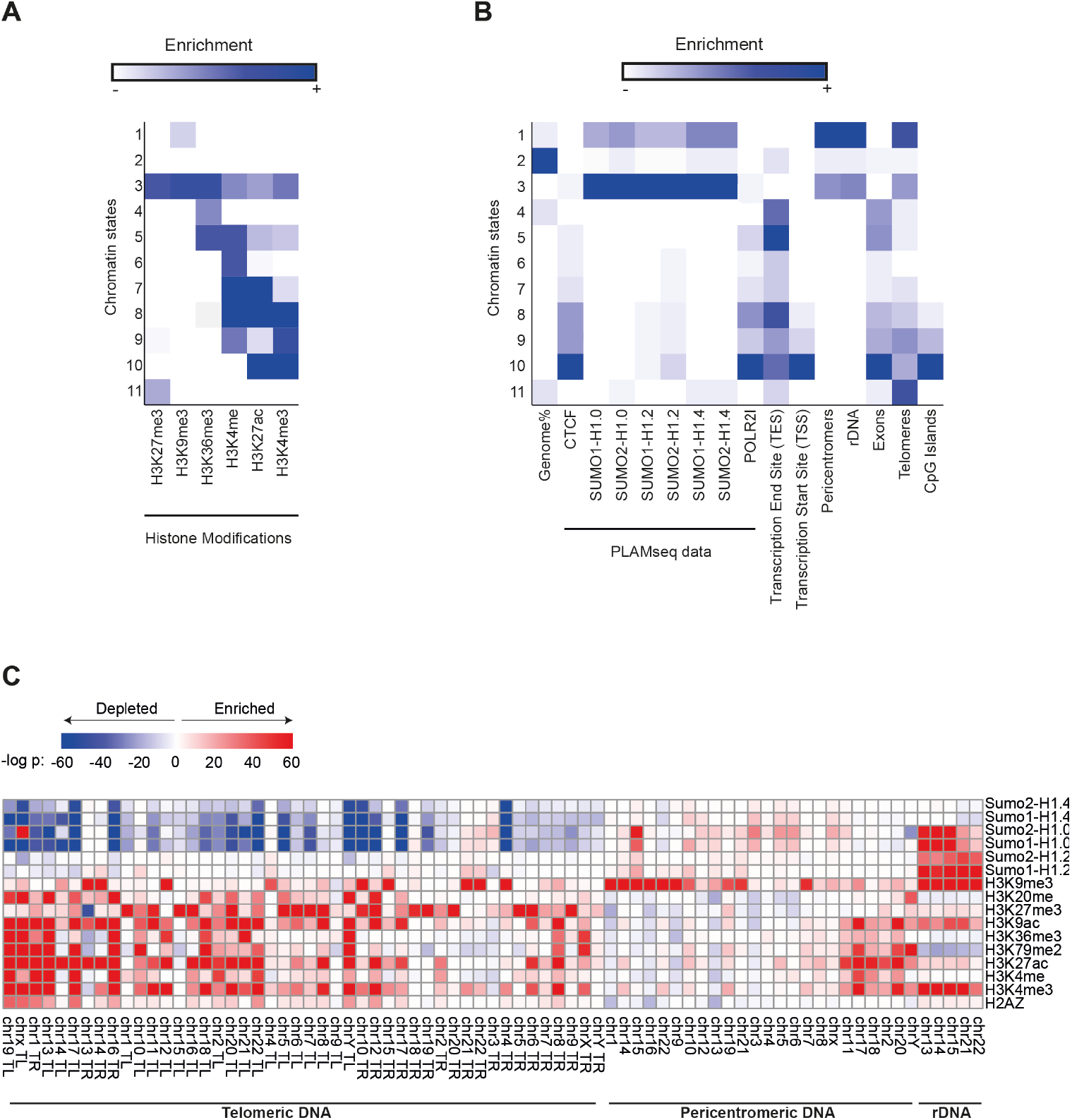
**(A)** ChromHMM heatmap showing the relative emission probability of each histone PTM for each of the 11 chromatin states in HeLa cells. **(B)**. Genome-wide enrichment of PLAMseq data and other genomic features across the different chromatin states. The columns represent the relative percentage of the genome covered by each chromatin state (first column) or the fold enrichment for the PLAMseq and the genomic annotations. **(C)**. Enrichment of histone PTMs (ChIP-seq) and sumoylation of different histone H1 variants (PLAM-seq) within telomeric, pericentromeric, and rDNA regions. The colour gradient corresponds to the -log10 of the FDR-adjusted p-value resulting from the comparison with the input (ChIP-seq) or the total H1 (PLAMseq), multiplied by the sign of the fold enrichment (see legend).

We found that SUMOylated histone H1 types accumulated at chromatin states in a similar manner as H3K9me3, as expected from the interaction with SETDB1. Moreover, repetitive regions such as Pericentromeres, ribosomal DNA (rDNA) and telomeres also presented similar distribution among chromatin states. Thus, we decided to look at specific repetitive genomic regions, such as the different rDNA loci, telomeres and pericentromeres (Figure 5C, Supplementary Figure 4). SUMOylated histones H1.2 and H1.0, but not H1.4 presented a high localization at rDNA, and the same was true for H3K9me3. Overall, all the SUMOylated histone H1 types presented a mild enrichment at pericentromeric regions. Regarding telomeres, most of them were depleted from SUMOylated histone H1.4 and H1.0, however, some of them presented a mild enrichment was observed in the telomeres of one of the arms in acrocentric chromosomes, also consistent with an accumulation of H3K9me3.

## DISCUSSION

### Advantages of PLAMseq

Here, we present PLAMseq, a new workflow that enables the simultaneous characterization of the genomic and proteomic environment of chromatin associated proteins. Moreover, we showed that PLAMseq provides a virtually equal output as ChIPseq experiments for CTCF and RNA polymerase II but with the advantage of being an antibody-free approach. This actually solves one of the bottlenecks for the performance of ChIPseq experiments and other antibody-based approaches such as CUT&RUN/TAG, as ChIP grade antibodies are often not available for the proteins of interest. Furthermore, antibodies are costly reagents which also face sensitivity and specificity issues.

Nevertheless, in cases where high quality ChIP-grade antibodies are available and financial resources are not a limitation for the research group, PLAMseq is still advantageous as it provides the opportunity of the identification of the proteomic environment of a proteins of interest in the same experimental protocol.

Another important limitation of ChIP-seq experiments is that the mapping of transient interactions is not possible; it is possible to map chromatin modifications but not chromatin modifiers. One of the advantages of TurboID is that it enables the identification of transient interactions. Thus, PLAMseq will enable the proteo-genomic characterization of chromatin modifiers. Moreover, it can allow to distinguish among different enzymes able to leave the same histone mark should be possible.

### PLAMseq for the genomic mapping of protein interactions

By using split-TurboID, protein interactors of the protein of interest can also be mapped in the genome plus their protein partners. Here, we combined it with SUMO-ID to study histone H1 SUMOylation, but it can also be applied for other protein complexes or to study interaction-dependent genomic distributions. It could be argued that this approach cannot distinguish, in the case of SUMO, whether the reconstitution of the biotinylating activity comes from the SUMOylation of the histone itself or from the SUMOylation of a histone interactor. However, specifically for SUMO, both cases are likely true, based on the group SUMOylation model proposed by the group of Stephan Jentsch ^41^ and the non-covalent SUMO proteome data showed that in more than 90% of the cases are also SUMOylation substrates ^35^. In any case, the determination of SUMOylation sites in the purified biotinylated proteins is possible by different workflows ^26-28,34,35,42^.

### Limitations of the PLAMseq approach

Our results for CTCF and RNA polymerase II PLAMseq studies (Figures 2-3) indicate that it is possible to substitute ChIPseq experiments for PLAMseq experiments with comparable or better results in a more affordable manner, as no antibody is required. Also, PLAMseq enables to perform these experiments for proteins which no ChIP-grade antibody is available. However, PLAMseq has also some limitations compared to ChIPseq or CUT&RUN/TAG. First, there are the limitations intrinsic to TurboID, as it is required to tag the desired protein with TurboID which might affect the functionality of the protein. While PLAMseq is ideal to profile ubiquitin(-like)-modified proteins, where Ubi-ID or SUMO-ID can be applied ^1^, it cannot be applied for other PTMs based on chemical modifications (i.e. phosphorylation, methylation, acetylation). Alternatively, a split-PLAMseq approach between the protein of interest and the modifying enzyme could be applied, but this alternative would be much more limited than using ChIPseq with a specific antibody (although this is not always available).

## METHODS

### Plasmid construction

To generate the inducible Gateway-TurboID destination vector, the HF-TULIP2 system ^43^ was employed to replace the ubiquitin sequence downstream of the Gateway^®^ cloning cassette with that of TurboID, using standard cloning techniques. Subsequently, LR Clonase reactions (Thermo Fisher Scientific) were carried out following manufacturer’s instructions, utilizing donor plasmids containing the cDNA of POLR2I and CTCF without a stop codon, with the Gateway-TurboID vector serving as the destination vector. Alternatively, plasmids encoding TurboID or NTurboID-H1(H1.2, H1.0 and H1.4) were constructed using standard cloning techniques, employing FLAG-TurboID-GSQ and FLAG-NTurboID-GSQ as vectors (Barroso-Gomila et al., 2021). For the split constructs of SUMO1 and SUMO2, site-directed mutagenesis was performed on MYCtag-CTurboID-GSQ-SUMO1nc or SUMO2nc (Barroso-Gomila et al., 2021) to remove the non-cleavable proline residue and concurrently substitute glutamine with arginine at positions 92 and 88 in SUMO1 and SUMO2, respectively. The modified sequences were then cloned into a constitutive lentiviral vector under the CMV promoter. After assembly, all vectors were validated by sequencing.

Additional details for constructs are described in Supplementary Table 1. Primer sequences are shown in Supplementary Table 2.

### Cell culture

HeLa and HEK 293T cells were cultured in Dulbecco’s modified Eagle Medium (DMEM) supplemented with 10% Fetal Bovine Serum (FBS) and Penicillin (100 U/mL)/Streptomycin (100 µg/mL) at 37°C and 5% CO2. Cells were tested negative for mycoplasma contamination.

### Lentiviral production

HEK 293T cells were seeded at 30% confluency in a T175 flask with 16 ml of DMEM with 10% FBS. The following day, a 2 ml transfection mixture was prepared in 150mM NaCl, containing lentiviral packaging plasmids: 7.5 μg of pMD2.G (Addgene, #12259), 11.4 μg of pMDLg-RRE (Addgene, #12251), 5.4 μg of pRSV-REV (Addgene, #12253), and 13.7 μg of lentiviral plasmids of TurboID or split-TurboID, along with 114 μl of Polyethylenimine (PEI, 1 mg/ml). After vortexing, mixes were incubated for 10 min at room temperature (RT) before being added to HEK293T cells. The day after transfection, culture medium was replaced with fresh DMEM + FBS/Pen/Strep. Three days post-transfection, the lentiviral suspension was collected and filtered through a 0.45-μm syringe filter (SLHVM33RS,Millex^®^-HV).

### Generation of stable cell lines

HeLa cells were seeded at 10% confluency in 15 cm plates with DMEM + 10% FBS. The following day, the culture medium was replaced with lentivirus-containing medium for either CTCF-TurboID, POL2RI-TurboID, TurboID-H1, NTurboID-H1, CTurboID-SUMO1, or CTurboID-SUMO2 constructs, along with polybrene (8 μg/ml). After 24 hours, the lentiviral medium was replaced with fresh DMEM + FBS/Pen/Step. Two days later, specific antibiotics were added to select for positive clones. Puromycin (3 μg/ml) was used to select CTCF-TurboID and POL2RI-TurboID cells, while a lower concentration (1 μg/ml) was applied for CTurboID-SUMO1 and CTurboID-SUMO2 cells. Blasticidin (5 μg/ml) was used for TurboID-H1.2 and NTurboID-H1.2 cells.

### Electrophoresis and immunoblotting

Input samples were lysed in 200-400 μl of SNTBS buffer (2% SDS, 1% NP-40, 50 mM tris pH 7.5, and 150 mM NaCl) and boiled at 100ºC. Samples were separated on Novex 4 to 12% gradient gels (Thermo Fisher Scientific) using NuPAGE Mops SDS running buffer (50 mM Mops, 50 mM tris base, 0.1% SDS, and 1 mM EDTA (pH 7.7)) or in-house casted gels of different Acrylamide percentages, proteins were transferred onto Amersham Protran Premium 0.45 NC nitrocellulose blotting membranes (GE Healthcare) using a Bolt Mini-Gel system (Thermo Fisher Scientific), which was used for both the gel electrophoresis and the protein transfer to the membrane according to vendor’s instructions. Membranes were stained with Ponceau-S (Sigma-Aldrich) to determine the total amount of protein loaded. Next, membranes were blocked with blocking solution (8% milk, 0.05% Tween-20 in PBS) for 1 hour before primary antibody incubation. Primary antibodies were incubated overnight at 4ºC and secondary antibodies for 1–2 hours at RT. Chemiluminescence reaction was initiated with Western Bright Quantum Western blotting detection kit (Advansta-Isogen) and measured in an ImageQuant 800 (Cytiva, Malborough, MA, USA) system. Antibodies are listed in Supplementary Table 3.

### PLAMseq (Proximity Labelled Chromatin Affinity Purification Mass Spectrometry plus sequencing)

A bench-top PLAMseq protocol is provided as Supplementary Protocol 1

### Biotin incubation and crosslinking

To perform PLAMseq, 60 million cells were seeded per experimental condition and treated with 50 µM biotin for 10 mins at 37°C. Cells were then washed with PBS, crosslinked with 1% formaldehyde in PBS and incubated for 20 mins at RT. Crosslinking was quenched by addition of 1.25 M glycine for 5 mins. Cell pellets were collected by centrifugation at 900 g for 5 mins, washed with PBS and stored at -80°C until use.

### Lysis, sonication and streptavidin pull-down

For cell lysis, each cell pellet was resuspended in 2 ml lysis buffer (50 mM HEPES pH 7.9, 140 mM NaCl, 1mM EDTA pH 8.0, 10% (vol/vol) glycerol, 0.5% (vol/vol) IGEPAL CA-630/NP-40 and 0.25% (vol/vol) Triton X-100) containing protease inhibitors and incubated for 10 mins on ice. Each sample was then split into two 1 ml aliquots and nuclei were collected by centrifugation at 1,700 g for 5 mins at 4°C. The supernatant was discarded, and the nuclei were resuspended in wash buffer (10 mM Tris-HCl pH 8.0, 1mM EDTA pH 8.0, 0.5mM EGTA pH 8.0 and 200 mM NaCl), incubated for 10 mins on ice, and centrifuged again. After rinsing the nuclei twice with shearing buffer (10 mM Tris-HCl pH 7.6, 1mM EDTA pH 8.0 and 0.1% (wt/vol) SDS), the nuclei pellet was resuspended in 1 ml of shearing buffer and transferred to AFA tubes. Chromatin was sheared using a Covaris ultrasonicator (PIP = 140, Duty = 5%, Time = 1020 sec, 6°C). After sonication, samples were centrifuged at 14,000 g for 10 mins at 4°C, and the supernatant was transferred to a new tube for the streptavidin pull-down. 25 µl of streptavidin beads (50% slurry) were added per sample and incubated overnight at 4°C with rotation. After incubation, beads were washed with shearing buffer and subjected to decrosslinking by incubating in decrosslinking buffer (1% SDS and 500mM NaCl in TE buffer 1X) at 65°C overnight shaking at 600 rpm.

### DNA purification and BioID washes

Following decrosslinking, samples were centrifuged at 500 x g and the supernatant (DNA) was diluted by half with TE buffer (10 mM Tris PH 8 and 1 mM EDTA) and incubated with 0.5mg/ml of proteinase K for 1h 30 mins at 37ºC. Samples were then further diluted by half with TE buffer to reduce SDS percentage and purified using a Gel and PCR clean-up kit (Macherey-Nagel).

Beads from the Streptavidin pull-down were washed sequentially with buffers 1 to 4 for 10 mins each (Buffer 1: 2% SDS; Buffer 2: 0.1% deoxycholate, 1% Triton X-100, 500mM NaCl, 1mM EDTA and 50mM HEPES pH 7.5; Buffer 3: 250 mM LiCl, 0.5% Triton X-100, 0.5% Deoxycholate and 1mM EDTA 10mM TrisCl pH 8.0 and Buffer 4: 50mM Tris HCl pH7.4 and 50mM NaCl). Beads were then washed three times with Ammonium bicarbonate (ABC) 50mM and finally resuspended in 250µl 50 mM ABC. Samples were then subjected to reduction with 1 mM DTT for 30 mins, alkylation with 5 mM chloroacetamide for 20 mins and a further incubation with 5mM DTT for 30 mins. Proteins were then digested overnight at 37°C using 250ng Trypsin, and peptides were separated from the beads using a 0.45µm centrifugal filter unit (Millipore).

### Library preparation

Library preparation for CTCF-TurboID, POL2RI-TurboID, and all split-TurboID/TurboIDH1.2 samples was conducted at the Genomic Unit of the Andalusian Center for Molecular Biology and Regenerative Medicine (CABIMER) (https://www.cabimer.es/unidades-apoyo/genomica/) using the ThruPLEX DNA-Seq kit (Takara, Cat. No. R400675).

For H1.4 and H1.0 samples, libraries were prepared using a tagmentation reaction with 10 µl DNA, 0.3µg Tn5 transposase and 6µl Tagmentation buffer 5x (50mM Tris pH 8.0, 25mM MgCl2, 50% v/v dimethylformamide) for 5 mins at 37°C. Reaction was stopped by addition of 3µl 1% SDS and samples were purified with 0.9x Sera-Mag beads (Cytiva) according to manufacturer’s instructions. Amplification was assessed by qPCR using 1 μl of the sample, 1X SYBR Gold (Thermo Fisher Scientific), 1X NEBNext® High-Fidelity PCR Master Mix, and 1 μl of Nextera DNA Indexes i5/i7 (https://support-docs.illumina.com/SHARE/AdapterSequences/Content/SHARE/AdapterSeq/Nextera/NexteraDNAIndexes.htm) at 10 μM. For library amplification and indexing of H1.4 and H1.0 samples, the NEBNext® High-Fidelity 2X PCR Master Mix was used with the following PCR program: 72°C for 5 mins; 98°C for 30 seconds; 9–10 cycles of 98°C for 10 seconds, 63°C for 30 seconds, and 72°C for 30 seconds; followed by a final extension at 72°C for 5 mins. Libraries were size-selected using dual selection with Sera-Mag beads at 0.7x-0.9x.

### Next Generation Sequencing

Libraries for all samples were sequenced at the Genomic Unit of CABIMER using the Illumina NovaSeq6000 SP-200 platform to generate 75 bp paired-end reads. Exceptions included input libraries and TurboID-H1.2 replicate 1, which were initially analyzed using the NextSeq500 MID-Output. All sequenced libraries were de-multiplexed, and adapter sequences were trimmed using the Illumina BCL Converter (version 2.20.0).

### Read Alignment and Preprocessing

Three independent replicates were performed for each PLAMseq experiment. Mapping and subsequent analysis of the data were done using Galaxy Server ^44^ (https://usegalaxy.eu) and custom R scripts. Reads were mapped to human genome assembly (GRCh38/hg38) using Bowtie ^45^ using parameters -m1--best. PCR duplicates were removed using ‘RmDup’ (Galaxy version 2.0.1) ^46^ with default parameters. For the analysis of histone sumoylation, raw sequencing reads were aligned to the telomere-to-telomere (T2T-CHM13v2.0) reference genome ^39^ using the Rsubread package ^47^ in R. The genome was first indexed using the buildindex function, and reads were aligned with the align function, allowing multi-mapping but retaining only the best-mapped location for each read. To minimize PCR amplification biases, duplicate reads were removed using samtools markdup -r ^48^. Only properly paired and non-duplicated reads were retained for downstream analyses. The resulting BAM files were sorted and indexed.

Genomic regions of interest were defined based on existing annotations. Position of genes and transcription start site (TSS) were obtained from Ensembl (Ensembl Genes 108). Transcriptionally active genes were selected as those having at least 10 FPKM in chromatin associated RNA experiments analyses previously published in Hela S3 ^49^ (GSM6787994). Silent genes were the rest. rDNA and pericentromeric regions were obtained from the UCSC Table Browser ^50^, while telomeres were defined as the 200 kb at the ends of each chromosome.

### Differential Enrichment Analysis

To quantify read enrichment across rDNA, pericentromeric, and telomeric regions, read counts were obtained using the windowCounts function from csaw ^51^. Sample normalization was performed with normFactors function using the TMM method and windows of 10 kb. Dispersion estimation was conducted using estimateDisp, and a negative binomial generalized linear model (GLM) was fitted using glmQLFit from edgeR^52^. Statistical significance was determined based on false discovery rates (FDR < 0.05). Results were visualized as a heatmap, where the represented value corresponds to the log10 of the FDR multiplied by the sign of the fold change. Volcano plots were generated using ggplot2 ^53^.

### Chromatin State Analysis

Publicly available histone modification datasets from the ENCODE project ^54^ (accession number ENCSR068MRQ) were processed to characterize chromatin states. Reads were binarized with ChromHMM ^40^. First, genome was segmented into 200-bp bins and binary values based on read presence were assigned using BinarizeBam function. Then, a chromatin state model with 11 states was trained using the LearnModel function in under default parameters. To assess chromatin state enrichment across genomic features, transcription start sites (TSS), transcription end sites (TES), and exons were extracted from the RefSeq annotation of CHM13 ^55^, while CpG island locations were retrieved from the UCSC Table Browser ^50^. Enriched regions for each histone H1 sumoylation and CTCF and POLR2I were identified by binarizing BAM files with BinarizeBed, using total histone samples as a control, and converting the binarized data to BED format. Enrichment of these regions was evaluated with the OverlapEnrichment function with default parameters.

### Peak Calling for CTCF and POLR2I

Peak calling of ENCODE CTCF sites was performed as above from the two BAM files available at GEO database (GSM733785). In this case, peaks present in both biological replicates were converted to (GRCh38/hg38) genome assembly using the LiftOver utility from UCSC genome browser ^56^ and used for the subsequent analyses.

### Signal Visualization and Coverage Analysis

Genome-wide coverage files were generated in BigWig format using bamCoverage from deepTools2 ^57^ with a bin size of 1 and normalized by CPM (counts per million). Then, the average signal from the three replicates were calculated using bigwigAverage from deepTools2 and used for the subsequent analyses. For CTCF, RNAPII, and firstly for histone H1.2, signal previously published by ENCODE (GSM733785 and GSM733759, respectively), coordinates were converted to (GRCh38/hg38) genome assembly using CrossMap ^58^. In all cases, signal distribution across the genome was analysed using UCSC genome Browser ^56^. Signal distribution across genomic features was analyzed using the computeMatrix function from deepTools2 to quantify signal intensity within genomic bins for each sample. For final profile metaplots and heatmaps, average signal of three biological replicates was computed and plotted. In heatmaps, genome positions were sorted from maximal to minimal intensity obtained in CTCF ENCODE data (GSM733785). Correlation heatmaps were done using multiBamSummary and PlotCorrelation tools from DeepTools using 1000bp non-overlapping bins.

### Mass spectrometry sample preparation

Digested peptides were acidified by adding 2% trifluoroacetic (TFA) acid. Subsequently, peptides were desalted and concentrated on triple-disc C18 Stage-tips as previously described ^59^. Stage-tips were in-house assembled using 200 µL micro pipet tips and C18 matrix (Sigma-Aldrich). Stage-tips were first activated by passing through 100 µl of methanol. Next, 100 µl of Buffer B (80% acetonitrile, 0.1% formic acid), 100 µl of Buffer A (0.1% formic acid), the acidified peptide sample, and two times 100 µl Buffer A were passed through the Stage-tip. Elution was performed twice with 30 µl of Elution buffer (32,5% acetonitrile, 0.1% formic acid solution). Samples were vacuum dried using a Universal Vacuum System UVS400S coupled to a SpeedVac SPD121P (Thermo) and stored at −20 °C. Prior to mass spectrometry analysis, samples were reconstituted in 20 µl 0.1% formic acid and transferred to autoload vials.

### Mass spectrometry data acquisition

Mass spectrometry-based proteomics data was acquired at the Proteomics facility in Institute of Biomedicine of Seville (IBiS). As previously done ^60^, samples were measured using a nanoElute II LC system coupled to a timsTOF SCP mass spectrometer with an electrospray source (Bruker Daltonics). LC separations were performed on C18 HPLC column (Aurora 25 cm and 75 µm ID, IonOpticks) kept at 50°C. Gradient elution was performed with a binary system consisting of (A) 0.1% aqueous formic acid and (B) 0.1% formic acid in Acetonitrile. An increasing linear gradient (v/v) was used (t (min), %B): (0, 2); (40, 17); (60, 25); (66, 37); (67, 95), followed by an equilibration step.

Mass spectrometric analysis was performed in a data independent acquisition parallel accumulation serial fragmentation (dia-PASEF) mode, with 100–1700 m/z mass range, an ion mobility range from 0.64 to 1.45 V s cm−2, capillary voltage set to 1500 V, an accumulation and ramp time at 100 ms and the collision energy as a linear ramp from 20 eV at 1/K0 = 0.6 V s cm−2 to 59 eV at 1/K0 = 1.6 V s cm^−2^.

### Mass spectrometry data analysis

Mass spectrometry raw dia-PASEF data was analyzed using Biognosys Spectronaut (v19.7.250203) using a directDIA+ search with BGS Factory settings. Predicted spectral library was built using a FASTA file corresponding to the reference human proteome (Uniprot 29^th^ August 2022). A pivot report was generated and further processed in the Perseus Computational Platform ^61^. Proteins not identified in every replicate for at least one condition were removed and missing values were randomly imputed using normally distributed values with 0.3 width and 1.8 down shift considering the total matrix. Subsequently, proteins which quantification had been performed based in only one (1) precursor were also removed. Statistical analysis was performed using two-sided Student’s t tests. Results were exported into in MS Excel 365 for a comprehensive browsing and visualization of the datasets. Volcano plots were constructed for data visualization using the VolcaNoseR web app ^62^ (https://huygens.science.uva.nl/VolcaNoseR2/).

## Supporting information

Spplemenatry Protocol 1

## DATA AVAILABILITY

The mass spectrometry proteomics data have been deposited to the ProteomeXchange Consortium via the PRIDE ^63^ partner repository with the dataset identifier PXD062203. The genomics data discussed in this publication have been deposited in NCBI’s Gene Expression Omnibus ^64^ and are accessible through GEO Series accession number GSE 294161.

## ACKNOWLEDGMENTS

Authors would like to thank Alfred Vertegaal, Jim Sutherland and Rosa Barrio for sharing plasmid constructs. Additionally, authors would like to thank Eloísa Andújar-Pulido, Mónica Pérez-Alegre, Victoria E. Jiménez and Lola P. Camino from the CABIMER Proteomics facility for the support and acquisition of genomics data. Also, would like to thank Gonzalo Millán-Zambrano for critical discussions about the PLAMseq method.

This work was supported by grants PID2021-122361NA-I00 by MICIU/AEI/10.13039/501100011033 and European Union, CNS2022-135216 funded by MICIU/AEI/10.13039/501100011033 and European Union NextGenerationEU/PRTR, the EMERGIA 2020 program (EMERGIA20_00276) from the Consejería de Economía, Conocimiento, Empresas y Universidad, Junta de Andalucía, Spain to RG-P. MES-O was hired by the QUALIFICA program of Junta de Andalucía.

## AUTHOR CONTRIBUTION

RG-P conceived the project. LG-V and CE-S experimental work. RG-P, LG-V, CE-S, MES-O, DR, and CG-A data analysis. MLM-M proteomics data acquisition. LG-V and CE-S were supervised by RG-P. MES-O was supervised by DR. RG-P wrote the manuscript with contributions from all authors.

