## Supplementary Figures and tables for "PLAMseq enables the proteo-genomic characterization of chromatin-associated proteins and protein interactions in a single experimental workflow"

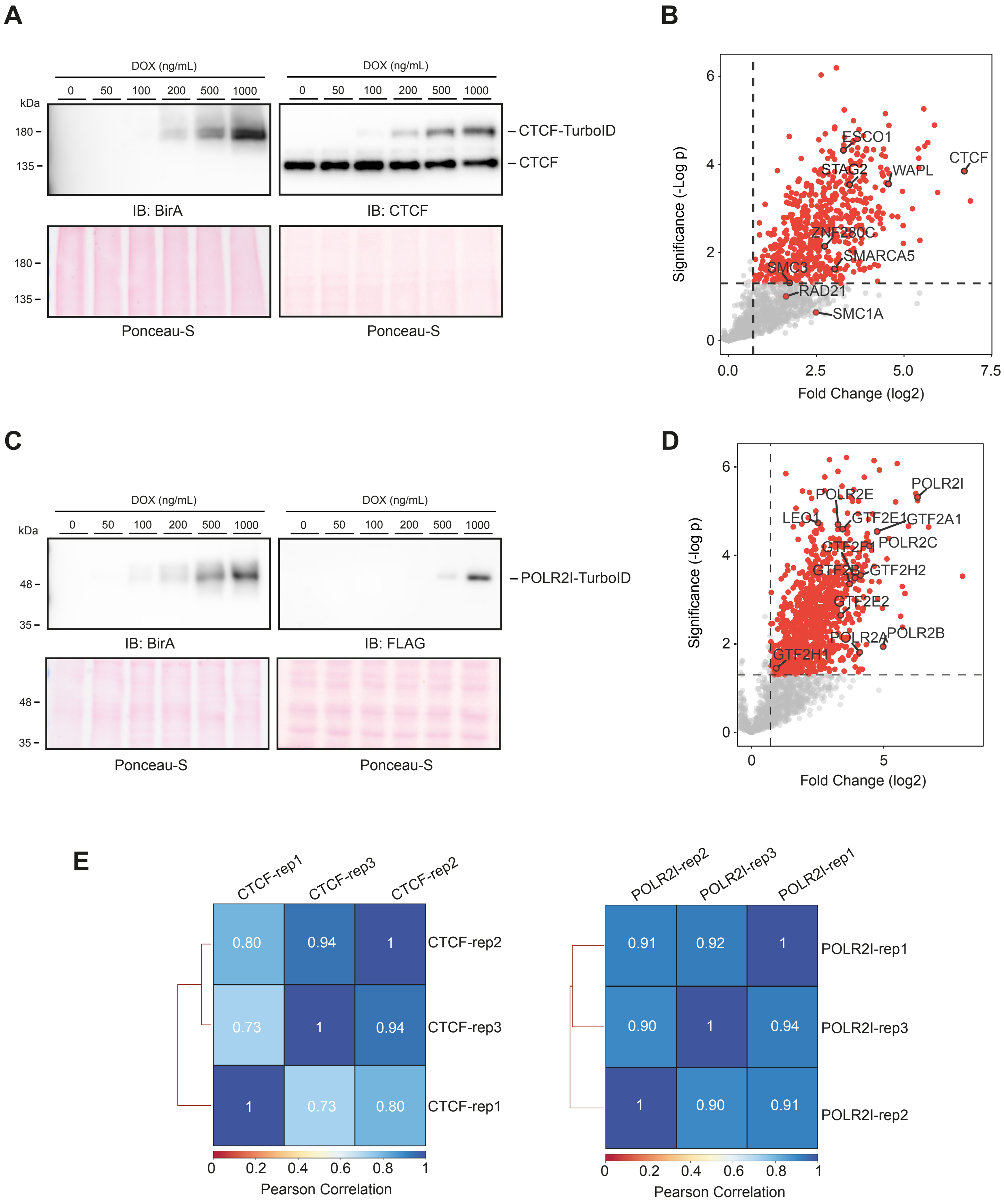


**Supplementary Figure 1: (A)** Immunoblot showing endogenous CTCF and CTCF-TurboID expression levels at different doxycycline concentrations. Ponceau Staining is provided as loading control. **(B)** Volcano plot depicting statistical differences of the proteomics analysis of CTCF-PLAMseq samples compared to Control samples. Each dot represents a protein. **(C)** Immunoblot of POLR2I-TurboID expression levels at different doxycycline concentrations. Ponceau Staining is provided as loading control. **(D)** Volcano plot depicting statistical differences of the proteomics analysis of CTCF-PLAMseq samples compared to Control samples. For the volcano plots (B,D) dotted lines limit significant threshold for a fold-change higher than 0.7 and a -log p higher than 1.3 (p>0.05) for unpaired two-tailed t-tests from three biological replicates.

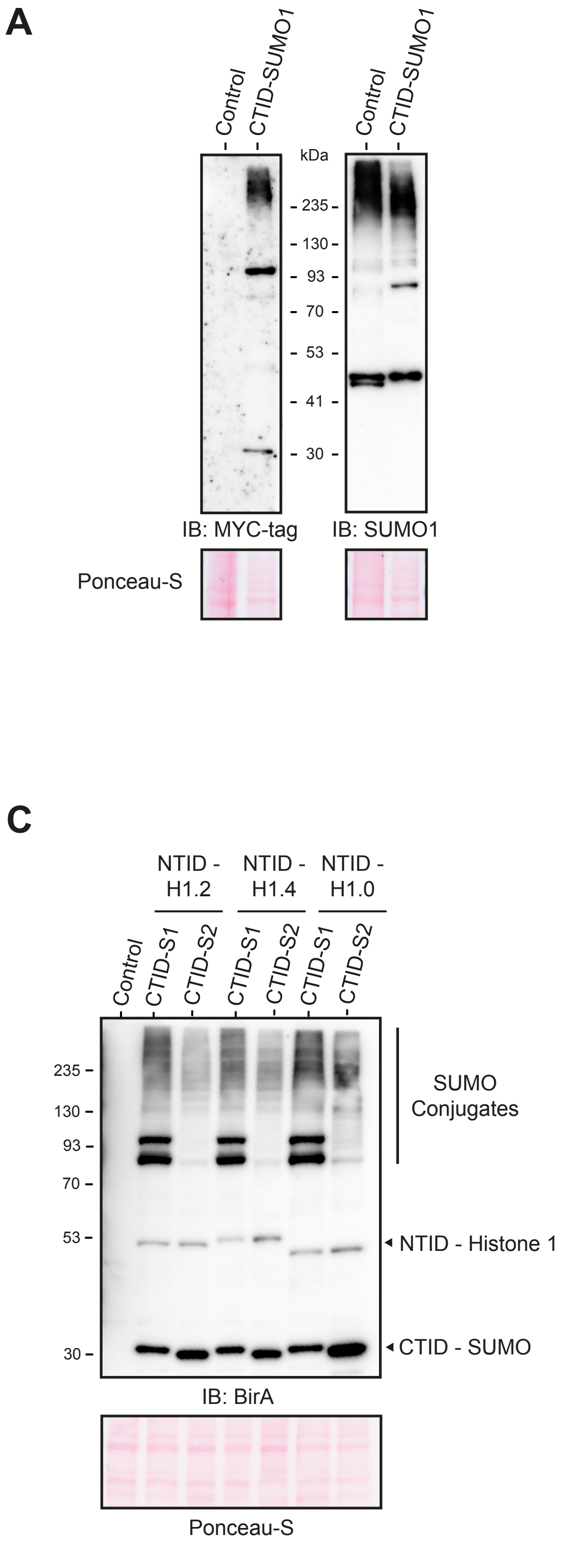


**Supplementary Figure 2: (A)** Immunoblots showing expression of myc-split-TurboID-SUMO1 and **(B)** myc-split-TurboID-SUMO2 in HeLa cells. **(C)** Immunoblots of HeLa stable cell lines for CTurboID-SUMO1 and CTID-SUMO2 in combination with NTurboID-Histone 1 isoforms H1.2, H1.4 and H1.0, and **(D)** for full lenght TurboID-H1 isoforms. Ponceau staining is provided as loading control. C-terminal TurboID and N-terminal TurboID are abbreviated as CTID and NTID, respectively.

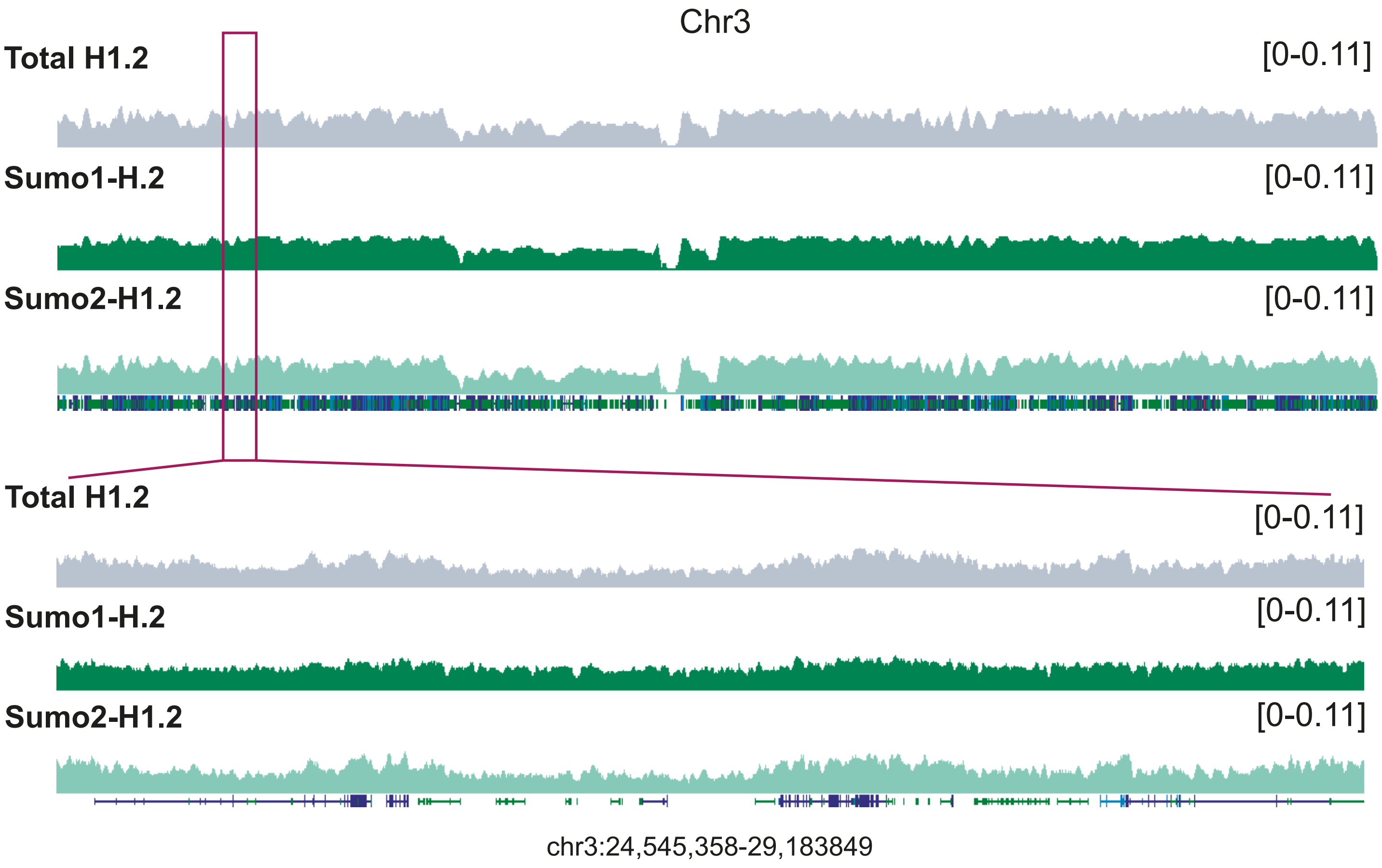


**Supplementary Figure 3.** Genomic profiles of Total Histone H1.2, SUMO1-modified Histone H1.2 and SUMO2-modified Histone H1.2

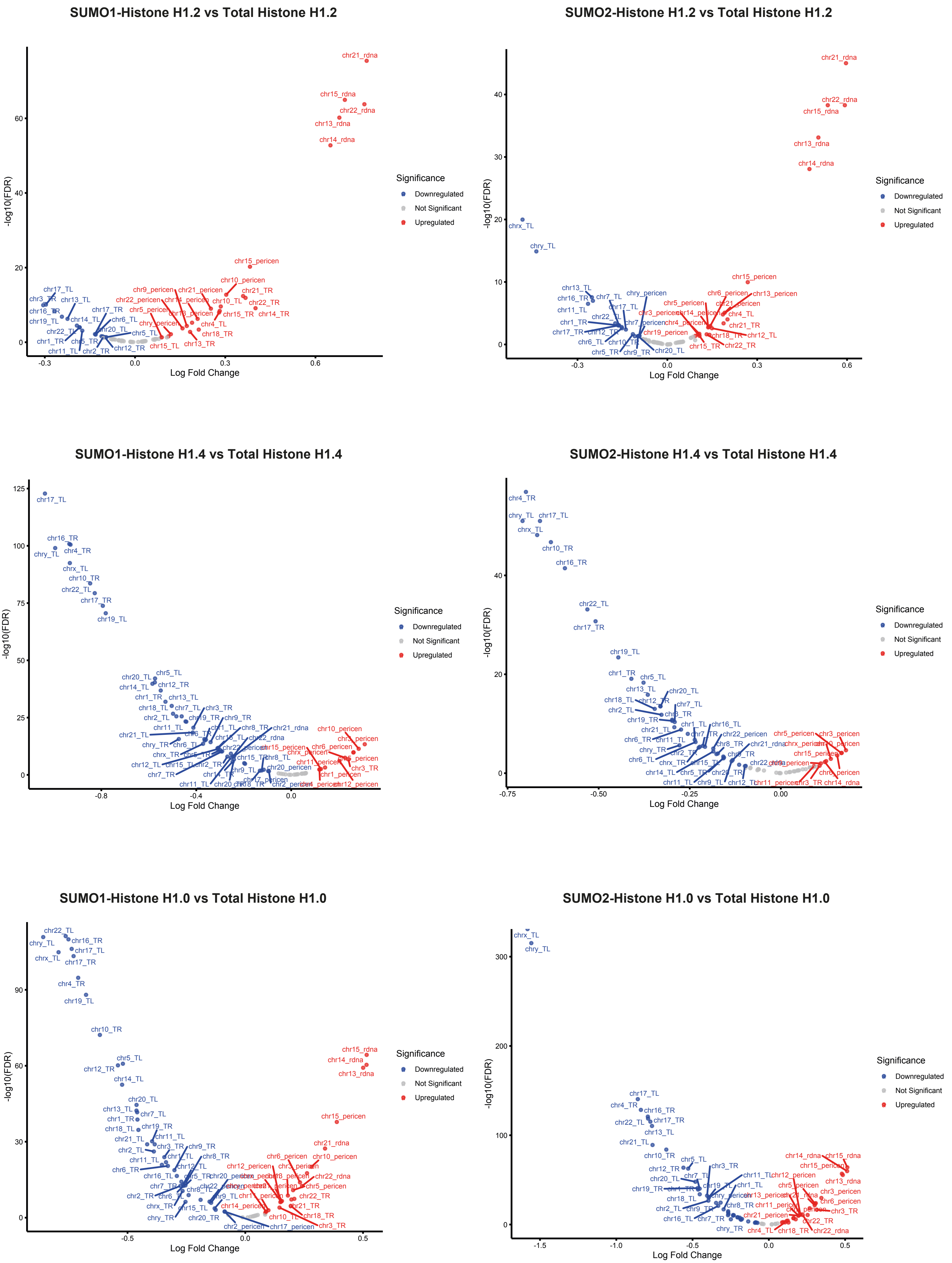


**Supplementary Figure 4:**Volcano plots depicting statistital enrichment at repetitive DNA genomic loci of SUMOylated Histone H1 types compared to their total Histone H1 type counterpart.

**Supplementary Information**

**TABLE 1. List of plasmids**

| Plasmid Number | Plasmid | Resistance |
|---|---|---|
| pRGP #405 | TREp-10xHis-FLAG-Gateway-GSQ-TurboID-PURO | AMP |
| pRGP #386 | DONR223-attL-POLR2I-attL | SPECT |
| pRGP #389 | DONR223-attL-CTCF-attL | SPECT |
| pRGP #482 | TREp-10xHis-FLAG-POLR2I-GSQ-TurboID-PURO | AMP |
| pRGP #483 | TREp-10xHis-FLAG-CTCF-GSQ-TurboID-PURO | AMP |
| pRGP #345 | EFSp-FLAG-TurboID-GSQ-H1.2-P2A-BLAST | AMP |
| pRGP #344 | EFSp-FLAG-NTurboID-GSQ-H1.2-P2A-BLAST | AMP |
| pRGP #356 | EFSp-FLAG-TurboID-GSQ-H1.4-P2A-BLAST | AMP |
| pRGP #355 | EFSp-FLAG-NTurboID-GSQ-H1.4-P2A-BLAST | AMP |
| pRGP #377 | EFSp-FLAG-TurboID-GSQ-H1.0-P2A-BLAST | AMP |
| pRGP #376 | EFSp-FLAG-NTurboID-GSQ-H1.0-P2A-BLAST | AMP |
| pRGP #403 | CMVp-MYC-CTurboID-GSQ-SUMO1-IRES-PURO | AMP |
| pRGP #404 | CMVp-MYC-CTurboID-GSQ-SUMO2-IRES-PURO | AMP |

**TABLE 2. List of primers**

| Primer Name | Sequence | Use |
|---|---|---|
| FW-EcoRI-H1.2 | GAATTCATGTCCGAGACTGCTCCTGC | Generation of TurboID/NTID-H1.2 construct |
| RV-BamHI-H1.2 | GGATCCTTTCTTCTTGGGCGCCGC | Generation of TurboID/NTID-H1.2 construct |
| FW-EcoRI-H1.4 | GAATTCATGTCCGAGACTGCGCCT | Generation of TurboID/NTID-H1.4 construct |
| RV-BamHI-H1.4 | GGATCCCTTTTTCTTGGCTGCCGCCT | Generation of TurboID/NTID-H1.4 construct |
| FW-MfeI-H1.0 | CAATTGATGACCGAGAATTCCACGTCCG | Generation of TurboID/NTID-H1.0 construct |
| RV-BamHI-H1.0 | GGATCCCTTCTTCTTGCCGGCCCTCT | Generation of TurboID/NTID-H1.0 construct |
| FW-NsiI-MYC-CTID | ATGCATATGGAACAAAAACTCATCTCAGAAGAG | Generation of CTID-SUMO1 and SUMO2/3 construct |
| RV-XbaI-SUMO1 | TCTAGATTAACCACCCGTAGGTTCCTG | Generation of CTID-SUMO1 construct |
| RV-XbaI-SUMO2/3 | TCTAGATTAACCTCCCGTTGGCTGTT | Generation of CTID-SUMO2 construct |
| FW-SUMO1 Q92R + P94Q | GGAGGAAGAAGATGTGATTGAAGTTTATAGGGAACAAACGGGTGGTTAATCTAGACCTG | Mutation of SUMO1 Q92R and Q95P |
| RV-SUMO1 Q92R + P94Q | CAGGTCTAGATTAACCACCCGTTTGTTCCCTATAAACTTCAATCACATCTTCTTCCTCC | Mutation of SUMO1 Q92R and Q95P |

| | | |
|---|---|---|
| FW-SUMO2/3 Q88R + P90Q | AGGATGAAGATACAATTGATGTGTTCCGACAGCAAACGGGAGGTTAATCTAGACCTG | Mutation of SUMO2/3 Q88R and Q90P |
| RV-SUMO2/3 Q88R + P90Q | CAGGTCTAGATTAACCTCCCGTTTGCTGTCGGAACACATCAATTGTATCTTCATCCT | Mutation of SUMO2/3 Q88R and Q90P |
| FW-AgeI-GSQ | ACCGGTGGCGGGGGATCTTCCGG | Generation of Gateway-TurboID |
| RV-SpeI-TurboID | ACTAGTTTACTTTTCGGCAGACCGCAG | Generation of Gateway-TurboID |

**TABLE 3. Antibodies**

| **Primary Antibodies** | | | | |
|---|---|---|---|---|
| **Antibody** | **Target** | **Dilution** | **Company** | **Catalog Number** |
| Rabbit anti-DYKDDDDK Tag (D6W5B) | Flag DYKDDDDK | 1:1000 | Cell Signaling Technology | 14793 |
| Mouse anti-CTCF (G-8) | CTCF | 1:500 | Santa Cruz | sc-271474 |
| Mouse anti-Myc-Tag (9B11) | Myc-tag | 1:1000 | Cell Signaling Technology | 2276S |
| Rabbit anti-SUMO1 | SUMO1 | 1:1000 | Cell Signaling Technology | 4930S |
| Mouse anti-SUMO 8A2 | SUMO2/3 | 1:1000 | 8A2, obtained From Developmental Studies Hybridoma Bank (DSHB), University of Iowa, in-house produced | |
| Rabbit anti-BirA | Recombinant E. coli BirA / Bifunctional BirA protein | 1:1000 | SinoBiological | 11582-T16 |
| **Secondary Antibodies** | | | | |
| **Antibody** | **Target** | **Dilution** | **Company** | **Catalog Number** |
| HRP-conjugated Goat-anti-mouse | Mouse IgG | 1:5000 | Jackson Immunoresearch | 115-035-146 |
| HRP-conjugated Goat-anti-rabbit | Rabbit IgG | 1:5000 | Jackson Immunoresearch | 111-035-003 |
