## Supplementary material for "PLAMseq enables the proteo-genomic characterization of chromatin-associated proteins and protein interactions in a single experimental workflow": Spplemenatry Protocol 1

### PLAMseq (Proximity Labelled Chromatin Affinity Purification Mass Spectrometry plus sequencing)

### TURBO-ID and split-TURBOID Proteins

#### REAGENTS:

- Biotin Stock Solution (20x): 1 mM biotin in ddH2O. Dissolve 12.2 mg biotin (Sigma #B4501) in 50 mL serum-free cell culture medium. Vortex and filter sterilize solution through 0.4 µM filter unit. Store at 4 °C for up to 8 weeks.
- Formaldehyde (10% (vol/vol); Ricca Chemical, cat. no. 3180-16).
- Glycine (Sigma Aldrich, cat. no. 410225).
- Biotin (Sigma-Aldrich, cat. no. B4501).
- **1x Lysis Buffer:** 50 mM HEPES pH 7.9, 140 mM NaCl, 1mM EDTA pH 8.0, 10% (vol/vol) glycerol, 0.5% (vol/vol) IGEPAL CA-630/NP-40 and 0.25% (vol/vol) Triton X-100.
- **1x Wash Buffer:** 10 mM Tris-HCl pH 8.0, 1mM EDTA pH 8.0, 0.5mM EGTA pH 8.0 and 200 mM NaCl.
- **1x Shearing Buffer:** 10 mM Tris-HCl pH 7.6, 1mM EDTA pH 8.0 and 0.1% (wt/vol) SDS.

  Add cOmplete Protease Inhibitor Cocktail (PIC) (Roche, cat. no. 11836170001) to a final concentration of 1X to each buffer before use.

- **1X TE buffer:** 10 mM Tris PH 8 and 1 mM EDTA.
- **Decrosslinking buffer:** 1% SDS and 500mM NaCl in TE buffer 1X.
- milliTUBE-1 mL with AFA fiber (Tube Ref. 520130 and Rack Ref. 500431)
- **Buffer 1:** 2% SDS.
- **Buffer 2:** 0.1% deoxycholate, 1% Triton X-100, 500mM NaCl, 1mM EDTA and 50mM HEPES pH 7.5.
- **Buffer 3:** 250 mM LiCl, 0.5% Triton X-100, 0.5% Deoxycholate and 1mM EDTA 10mM TrisCl pH 8.0.
- **Buffer 4:** 50mM Tris HCl pH7.4 and 50mM NaCl.
- **5X tagmentase buffer:** 50mM Tris pH 8.0, 25mM MgCl2, 50% v/v dimethylformamide.

#### METHODS:

##### Biotin incubation and crosslinking

- Begin with 60 million cells for each experimental condition.
- Change medium to fresh complete medium containing 50µM biotin to activate Biotinylation. Incubate for 10 mins at 37 °C in a cell culture incubator.
- Crosslinking:
  1. Remove media and wash once with cold PBS 1X.
  2. Fix the cells by adding 10 ml of 1% (wt/vol) formaldehyde in PBS and incubating for 20 mins at room temperature (RT).
  3. Quench crosslinking by adding 1 ml of 1.25 M glycine 5 mins RT.
  4. Wash once with cold PBS 1X.

5. Add 3 ml of cold PBS 1X and collect the sample by scraping with a cell scraper. Transfer it to a 50 ml tube.
6. Centrifuge for 5 mins at 900 g at 4°C. Decant the supernatant.
7. Wash the cell pellet with 15 ml of cold PBS 1X.
8. Centrifuge for 5 mins at 900 g at 4°C. Decant the supernatant.
9. Cell pellets can be snap-frozen at this point and stored at -80 °C.

#### Lysis, sonication and streptavidin pull-down

1. If cells were frozen after formaldehyde fixation, thaw cells on ice first. Add 2 mL Lysis Buffer containing 1× protease inhibitors to cross-linked cells (60M) to lyse the plasma membrane, gently resuspend by aspirating/dispensing 4 times or until homogeneous.
2. Divide the sample into two Eppendorf tubes (1ml each). The following steps will be carried out considering each half of the sample (30M) as an individual sample.
3. Incubate for 10 mins on a rocker at 4°C.
4. Collect intact nuclei by centrifugation at 1,700 g for 5 mins at 4 °C. Decant the supernatant without disturbing the nuclei pellet.
5. Gently resuspend pellet (30M each) in 1 mL Wash Buffer containing 1x protease inhibitor and incubate on a rocker for 10 mins at 4 °C.
6. Collect nuclei by centrifugation at 1,700 g for 5 mins at 4°C. Carefully remove and discard the wash solution, taking care not to disturb the nuclei pellet.
7. Gently rinse the sides of the tube with 1 mL Shearing Buffer containing 1x Protease inhibitor. Slowly dispense the buffer down the entire circumference of the upper-inside of the tube, taking care not to disturb the nuclei pellet.
8. Collect nuclei by centrifugation at 1,700 g for 5 mins at 4 °C. Decant the supernatant without disturbing the nuclei pellet.
9. Repeat steps 7 and 8 an additional time. Carefully remove and discard the supernatant, taking care not to disturb the nuclei pellet.
10. Resuspend nuclei pellet (30M) in 1 mL Shearing Buffer and transfer to appropriate AFA Tube.
11. Shear chromatin with an AFA Focused-ultrasonicator (Covaris) with appropriate rack or holder and the following settings: PIP = 140, Duty factor = 5 %, CPB = 200, Time = 1020 seconds, Temperature = 6 °C (min. 3 °C – max. 9 °C), AFA Intensifier = Yes and Water level = 5.
12. Centrifuge sonicated samples at 14,000 g 10 mins 4 °C.
13. Transfer supernatant to a new Eppendorf.
    - Take 20 µL of sample as a sonication control.
    - Add 80ul of TE 1X buffer and RNAse A 50ug/mL (0,5ul of 10mg/mL stock). Incubate for 30 minutes at 37°C. Add 0,5% SDS and 0,5mg/ml of proteinase K and incubate 10h at 37 °C and a further 6h at 65 °C in a thermocycler.
    - Take 50ul as input.
14. Use 25µl streptavidin beads (50µl of slurry) per sample, previously equilibrated with shearing buffer. Incubate samples with beads at 4 °C O/N with rotation.

#### Sonication control and decrosslinking

- After overnight decrosslinking of the sonication control, dilute samples to half with TE 1X and purify DNA with a Gel and PCR clean-up kit. Run purified samples on 0,8% agarose gel (fragments are at 100-500bp).

- For the samples of the streptavidin pull-down, centrifuge them at 500 g for 3 mins and discard supernatant. Wash beads twice with 1ml shearing buffer + PIC at 4 °C, 5-10 mins on rotation. Add 200µl of decrosslinking buffer to beads and 150µl to input samples and leave samples at 65 °C 600 rpm overnight.

#### DNA purification and BioID washes

A. Centrifuge beads 500 g 3mins. Take supernatant (DNA) to a new Eppendorf (200ul).
   1. Add 200µl of TE 1X to inputs and PLAMseq DNA samples and 0.5mg/ml of proteinase K. Incubate 1h 30 mins 37 °C 600 rpm.
   2. Add 200µl TE 1X buffer and purify DNA with a Gel and PCR clean-up kit (elute in 20µl). Store -20 °C.

B. Wash beads (1ml each) with:
   1. Wash 1x 10 mins at RT with 1ml Buffer 1 on a rotator.
   2. Wash 1x 10 mins at RT with 1ml Buffer 2 on a rotator moving to LoBind tubes.
   3. Wash 1x 10 mins at RT with 1ml Buffer 3 on a rotator.
   4. Wash 1x 10 mins at RT with 1ml Buffer 4 on a rotator moving to LoBind tubes.
   5. Resuspend beads in 250µl Buffer 4 and save 10% of beads for western blot analysis.
   6. Discard remaining Buffer 4 and perform 3x washes with freshly prepared 1ml Ammonium bicarbonate (ABC) 50mM. Change to LoBind tubes with the second wash.
   7. Resuspend beads in 250µl 50mM ABC.
   8. Reduction-alkylation:
      - Add 1mM DTT for 30 mins at RT on a rotator.
      - Add chloroacetamide 5mM for 20 mins at RT on a rotator.
      - Add 5mM DTT for 30 mins at RT on a rotator.
   9. Add 250-500ng Trypsin.
   10. Incubate O/N at 37 °C 1,300 rpm.
   11. Separate the beads from peptides with a prewashed 0,4um centrifugal filter unit (Millipore).

#### Desalting by Stage-Tip and Mass Spectrometry preparation

1. Wash 0.45 µm filter columns with 200 µL of freshly prepared ABC buffer and centrifuge for 1 min at 8,000 g.
2. After digestion, slowly spin down the samples at 8,000 g for 1 min in the pre-washed 0.45 µm filter column, collect the digested proteins in a LoBind eppendorf tube and remove the filter with beads.
3. If Stage-Tip is not performed immediately, snap freeze the digested peptides (in the LoBind tubes) and storage them at -80 ºC.
4. Desalt peptides according to the Stage-Tip protocol (Rappsilber et al., 2007):
   - Acidify your peptide sample by adding trifluoroacetic acid (TFA) to 2%.
   - Take perforated tubes with C18 tips and activate/equilibrate them as follows:
     1. Add 100µL of MetOH. Centrifuge for 20-40 seconds at 1,000 g. Always watch the flow of the solution and do not let the filter of C18 tips to dry out.
     2. Wash with 100µL of 80% ACN, 0.1% FA. Centrifuge at 1000 g.
     3. Equilibrate the StageTips with 100µL 0.1% FA.

4. Load your sample over the StageTips. This may be done in multiple steps (max. 100µL).
5. Wash twice the StageTips with 100µL 0.1% FA. Centrifuge for 5-10 mins at 1,000 g.
6. Prepare regular 1.5 mL tubes by using a pair of scissors to puncture a small hole in the lid. Place the washed StageTips into the 1.5 mL tubes.
7. Perform the elution from the StageTips with 33% Acetonitrile, 0.1% Formic Acid. Elute twice with 30µL, centrifuging for 3 mins at 1000 g (total volume 60 uL).
8. Lyophilize peptides using a Universal Vacuum System UVS400S coupled to a SpeedVac SPD121P (Thermo). Once lyophilized (samples should be completely dry), keep them at -20 °C if Mass-Spectrometry analysis is not performed immediately.
9. Resuspend each sample in 20µL 0.1% FA.
10. Place them in an ultrasonic bath for two minutes.
11. Spin down samples and transfer to autoload vials and keep them at -20 °C until analyzed by mass-spectrometry.

#### Tagmentation

1. Bring Seramag Beads (Cytiva) to RT (at least 30 mins) and vortex to fully resuspend beads.
2. Prepare 30µl Tagmentation reaction (10µl DNA, 0.3ug Tn5 transposase, 6µl Tagmentation Buffer 5X and 12µl H2O) and incubate for 5 mins at 37 °C.
3. Stop reaction by adding 3µl 1% SDS.
4. Purify with 0,9X Seramag beads. Mix and incubate for 5 mins at RT.
5. Place samples on a magnet rack and incubate for 5 mins at RT.
6. Remove and discard clear supernatant taking care not to disturb the beads.
7. Wash 2X with 200µl EtOH 70%. With the tubes on the magnet, add 200µL of freshly prepared 70% ethanol to each magnetic bead pellet and incubate at RT for at least 30 seconds. Carefully, remove ethanol by pipette.
8. Remove tubes from the magnetic stand and let dry at RT for 5 mins or until dry. Don't over-dry beads.
9. Resuspend in 22µl miliQ water. Mix thoroughly by pipetting.
10. Incubate resuspended beads at RT for 5 mins.
11. Place tubes on the magnet at RT for 5 mins or until the supernatant appears clear.
12. Transfer 20µl of clear sample to a new tube.

#### Library preparation

1. Check number of cycles by qPCR (Vf = 20ul)
   - 1µl of 10µM primer mix Ni5/Ni7
   - 1µl DNA
   - 0.2µl SyBR Gold 100x (Thermo Fisher Scientific)
   - 10µl NEBNext High-Fidelity 2X PCR Master Mix
   - 7.8µl $H_2O$ miliQ

   Program in PCR:

   - 72 °C 5 mins

- 98 °C 30s
- 98 °C 10s
- 63 °C 30s — X40 cycles.
- 72 °C 30s
- 72 °C 5 mins

Select the number of cycles to reach 1/3 of the maximum fluorescence (Approx 1M).

2. PCR for library amplification (same program as before, but changing the number of cycles for each sample):
   - 1µl of 10µM primer mix Ni5/Ni7. Change i7 primer for each sample.
   - 10µl DNA
   - 14µl miliQ water
   - 25µl 2X Nextera PCR mix

#### Library size-selection

1. Put Seramag beads at RT 30 mins before starting the procedure.
2. Vortex Seramag beads and add 0.7x volumes of beads (35µl to 50µl volume of sample). Mix and incubate at RT for 5 mins.
3. Place tubes on the magnet at RT for 5 mins or until supernatant appears clear.
4. Transfer 80µl of clear sample to a new tube. Be careful not to disrupt the magnetic bead pellet or transfer any magnetic beads with the sample.
5. Add 0.15x volumes of Seramag beads (12µl to 80µl volume of sample). Incubate at RT for 5 mins.
6. Place tubes on the magnet at RT for 5 mins or until the supernatant appears clear.
7. Remove and discard clear supernatant taking care not to disturb beads.
8. With the tubes on the magnet, add 200ul of freshly prepared 70% EtOH to each magnetic bead pellet and incubate at RT for 30 seconds. Carefully, remove EtOH by pipetting.
9. Repeat for a total of two EtOH washes. Ensure all ethanol has been removed.
10. Remove tubes from the magnet and let them dry at RT for 5 mins.
11. Resuspend dried beads with 22µl of H2OmQ. Mix thoroughly by pipetting. Ensure beads are no longer attached to the side of the tube.
12. Incubate resuspended beads at RT for 2-5 mins.
13. Place tubes on the magnet at RT for 5 mins or until the supernatant appears clear.
14. Gently transfer 20µl of clear sample to a new tube.
